# PAPET: a collection of performant algorithms to identify 5-methyl cytosine from PacBio SequelII data

**DOI:** 10.1101/2023.03.17.533149

**Authors:** Groux Romain, Xenarios Ioannis, Schmid-Siegert Emanuel

## Abstract

**Background:** Cytosine followed by guanine (CpG) rich genomic regions are known important regulatory sequences among vertebrates. Their cytosines can be subjected to a variety of chemical modifications, one of which being 5-methylcytosine (5mC), usually coined CpG methylation. CpG methylation has been demonstrated to have a deep impact on the nearby regulated genes. The advent of single molecule real time sequencing methods have proven to be powerful enablers in this field of research because of the possibility to detect DNA chemical modification directly from the raw sequencing data. In this perspective, several 5mC detection methods have been proposed.

**Results:** In this work we present PAPET - PacBio Prediction of Epigenetic Toolkit - a computational method to detect 5mC directly from PacBio SequelII raw sequencing data. We characterized the 5mC signatures in the raw data and proposed a framework to model them. We also assessed the effect of the DNA sequence alone on the signal and propose a normalization method to leverage this effect. From this, we designed a probabilistic approach to predict the presence of cytosine methylation from the sequencing kinetics and benchmarked it.

**Conclusions:** We apply this framework to predict CpG methylation from SequelII data and demonstrate that the classifiers compare equally with PacBio’s prediction method counterparts in terms of AUC and achieve very high specificity and precision.

## 1 Background

The genomes of vertebrates are interspersed with cytosine followed by guanine (CpG) dinucleotides. CpGs are palyndromic sequences, meaning that CpGs always occur by pairs on the opposite DNA strands. CpGs can appear isolated in the genome or as part of CpG rich regions. The latter regions are termed CpG islands (CGI). CGIs are genomic regions with more CpGs than expected by chance and account for ∼ 5% of all CpGs in vertebrate genomes. The remaining ∼95% of CpGs occur individually in the genome [1].

CpGs can undergo a variety of covalent modifications [2], one of them being the deposition of a methyl group on the 5th atom of the cytosine ring, resulting in the creation of 5-methylcytosine (5mC) [3]. This phenomena is termed CpG methylation. CpG methylation is catalyzed by a dedicate set of DNA-methyl-transferases (DNMT). In human, the factor DNMT1 and the DNMT3 protein family are responsible for methyl deposition. DNMT1 shows a strong specificity for hemi-methylated CpGs and ensures that both strands of the DNA share the same methylation status whilst the DNMT3 family members are responsible for de novo CpG methylation [2]. Demethylation, on the other hand, is performed through DNA replication and oxydation of 5mC into 5-hydroxymethylcytosine (5hmC) and 5-formylcytosine (5fC) by Tet factors [4].

Interestingly, individual CpGs are largely methylated in vertebrate genomes while CGIs remain overwhelmingly unmethylated [5]. This observation has been linked with the fact that vertebrate genomes contain less CpG dinucleotides than expected by chance. From this, the following hypothesis has been proposed to explain the CGI landscape in vertebrate genomes : throughout evolution, transitions are ”cleaning” vetebrate genomes from individual methylated CpG, leaving behind CpG rich unmethylated regions [6].

Furthermore, evidences have suggested that CGIs are important functional genomic sites. It is estimated that ∼70% of vertebrate gene promoters are associated with a CGI [7]. In human and mouse, ∼50% of all CGIs fall close to a transcription start site (TSS) [8]. These observations not only highlight a strong link between CGI and transcriptional regulation but makes of CGI the most important promoter architecture of vertebreate genomes [7, 9].

The latest years of research have shed light on the mechanism through which CGI work to recruit the transcriptional machinery. Unmethylated CGIs seem capable to specifically recruit ZF-CxxC DNA binding domain proteins [10]. In turn, these factors enforce a loose chromatin structure through the recruitement of chromatin remodellers and histone methylases and histone demethylases. Eventually, CGIs are made H3 lysine 4 tri-methylation (H3K4me3) rich and H3 lysine 36 di-methylation (H3K36me2) depleted [10]. Additionally, CGI sequences seem to be inheritently nucleosome repulsing compared to other genomic region [8]. These conditions set a favorable environment for the recruitement of the RNA polymerase II (RNAPII) complex [10], that subsequently spontaneously engages in non-productive transcription of short RNAs [8]. The activation of the downstream genes only requires the recruitement of activating TFs to engage the RNAPII into productive/elongating transcription [10]. This strongly supports a model in which unmethylated CGIs allow to create a transcriptionally permissive chromatin environment. These observations suggest that, at rest, CGIs are readily responsive promoters upon activation signaling.

On the other hand, CGIs have been demonstrated to be able to recruit poly-comb complex members such as PRC2. This triggers the deposition of the repressive H3 lysin 27 tri-methyl (H3K27me3) histone mark. Together with H3K4me3, this leads to the creation of ”bivalent” (or poised) promoters [10]. Bivalent promoters seem to temporarily repress genes that will be needed later in development [10]. Finally, a few number of CGI promoters seem to be constitutively silenced. The exact mechanisms through which this happens are not yet clear. However it seems that it required the action of poly-comb members, the deposition of H3K27me3 and the methylation of the CGI [8, 10]. Eventually, this leads to the formation of hetero-chromatin and enforces a stable gene silencing

The role of individual CpGs in regulatory processes shall not be overlooked, even if less well studies. It has been demonstrated that many TF actually recognize CpG. This allows them to bind CGIs but does not limit them to this. Recent work has demonstrated that CpG methylation modulates the affinity of TFs for their binding sites, here again pointing to a functional role of CpG methylation in gene regulation [11].

In terms of genomic sequencing, whole genome bi-sulfite sequencing (WGBS) has been considered the gold standard to assess genome-wide CpG methylation for many years. This protocol has been adopted by major international efforts such as the ENCODE Consortium [12], the NIH Roadmap Epigenomics Consortium [13], the Blueprint Project [14] or the IHEC [15]. WGBS detection of 5mC relies on the conversion of unmethylated cytosine into uracil, leaving 5mC as is in the library. Thus any CpG found in the reads can be considered methylated. However, WGBS cannot differentiate between 5mC and other cytosine methylations forms such as 5hmC, creating dead angles in the field of epigenetics. Additionally, WGBS requires more library preparation steps than e.g. whole genome sequencing (WGS).

The emergence of single molecule real-time sequencing (SMRT-seq) sequencing technologies has been a game changer in this regard. Oxford Nanopore Technologies (ONT) and Pacific Biosciences (PacBio) have revolutionize the field of genomics, and epignetics is no exception to this rule. Both of these SMRT-seq technologies leverage a real-time monitoring of events at the single DNA molecule level [16, 17]. This leads to the creation of so-called kinetic data for each DNA molecule being sequenced. Decisive for epigenetic studies, chemical DNA modification have been demonstrated early on to impact the sequencing kinetics, enabling the detection of DNA chemical modification directly from WGS [18–20]. In the case of PacBio sequencing data, the kinetic data are readily accessible from the raw sequencing data and are encoded into two distinct variables : pulse width (PWD) and interpulse duration (IPD). PWD refers to the duration of a fluorophore signal pulse and IPD to the time elasping bewteen two such pulses. The obvious advantages of studying 5mC using long read technologies are that i) different types of CpG methylation can be differenciated and ii) no dedicated library preparation step is required, unlike WGBS. This opportunity lead to the development of dedicated computational methods to detect CpG methylation from SMRT sequencing data.

For ONT data, both supervised and unsupervised approaches have been explored. Nanoraw [21] is an unsupervised approach that uses a Mann-Whitney testing to identify positions in the genome with sequencing kinetics differing from a whole genome amplified (WGA) control and then cluster them to isolate plausible DNA modification signatures. It does not stricto sensu identify a DNA modification, it rather detects regions of divergence from the control in the genome, without telling the causing DNA modification. As supervised approaches, Nanopolish [22] implements a Hidden Markov Model (HMM) approach to call bases from the raw signal. The HMM was extended to call 5mC, in addition to putative DNA bases. Different deep learning approaches have also been explored. Megalodon [23] uses a Recurrent Neural Network (RNN) and has been released by ONT to call bases and can identify 5mC. DeepSignal [24] can identify 5mC and 6mA and relies on a 3-tier method that uses a Convolutional Neural Network (CNN) to construct a feature matrix from the raw kinetics that is then fed into a bidirectional RNN and finally into fully connected neural network to compute the final predictions. DeepMod [25] uses a bidirectional RNN with long short-term memory (LSTM) mechanism. Finally, METEOR [26] is a consensus approach that computes predictions using Megalodon and DeepSignal and that uses either a random forest or a multiple linear regression to achieve a final decision. Concerning PacBio AgIn [27] has been developed for RSII derived sequencing data. It extracts IPDs in a window of 21bp around the CpG of interest. Because the CpG methylation is overwhelmingly symetrical among between strands, AgIn pairs the kinetics on both strands together between matching positions. The IPDs are then normalized to account for the DNA sequence, resulting in a feature vector for the CpG of interest. Then a Linear Discriminant Analysis (LDA) is used to decide whether the CpG is methylated or not. These predictions are then input into a segmentation method that leverage the uncertainty of individual CpG predictions by accounting for what happens in the neighbouring CpGs, like in CGIs. The HK model [28] was developed for newer models of PacBio sequencers, namely the SequelI and SequelII. It constructs a feature matrix using the Circular Consensus Sequence (CCS) sequence, IPD and PWD kinetics from both strands. The opposite strand kinetics are matched together similar to AgIn. The feature matrix is then put in a CNN to predict the presence of 5mC on each read. Event hough the HK method achieves a very good prediction, its source code and models were not made readily available, limiting its usefulness for the research community. Similarily, PacBio released primerose, a CNN-based method, to compute per read methylation prediction. These predictions can then be feed in pb-CpG-tool [29] - another PacBio software - to compute genomic CpG predictions. Like AgIn, pb-CpG-tool accounts for neighbouring CpG status when predicting a CpG methylation. Finally, ccsmeth [30] uses a RNN with Bidirectional Gated Recurrent Unit (BiGRU) and attention mechanism to compute a per read methylation probability from two 21bp long feature matrices. Each feature matrix is constructed from one strand of the read and contains the read sequence and the corresponding IPD and PWD kinetics. Both matrices are given to the input layer and are later on connected in the RNN. The reads are then mapped against a reference and the individual read methylation predictions are fed in another RNN to predict the methylation status of a given CpG. Similar to AgIn and pb-CpG-tools, the second RNN accounts for the neighbouring CpGs of the CpG of interest in the decision process.

Most of the recent predictor methods rely on deep neural networks achieving impressive prediction power. However, deep learning approaches are well known to be data hungry when it comes to training and computationally intensive. Additionally, some of the recent methods and/or models are not readily accessible, e.g. the HK method [28] and, PacBio’s primerose. This situation has motivated us to design a linear time algorithm with lightweight models to perform CpG methylation prediction. In this paper, we describe the ”PacBio Predicting Epigenetics Toolkit” (PAPET) algorithms. PAPET is built around a mixture model framework to leverage PacBio kinetic signal, train models and use them as predictors to detect CpG methylation using PacBio sequencing data and thereby offering the ability of creating tailored models and predictors.

## 2 Results

To investigate the effect of CpG methylation on the kinetic signal, we used a SequelII dataset generated by PacBio. This dataset originates from the HG002/NA24385 individual. The dataset is composed of two samples: i) a negative sample subjected to WGA - which removed any chemical modification from the DNA - and ii) a positive sample derived from the negative sample that was subsequently subjected to SssI enzymatic treatment. SssI is a DNA methylase that specifically targets CpG for 5mC deposition. Both samples were then subjected to sequencing using a PacBio SequelII machine.

### 2.1 5mC kinetic signatures

We verified whether a difference of kinetic signal could be detected between the WGA and SssI samples, around CpGs. We opted for a reference based approach and mapped the CCSs against the hg38p8 genome. Suzuki and colleagues [27] have demonstrated that the 5mC kinetic signatures can be detected using 21bp large windows centered on the CpG. We extended this window to 31bp and extracted the IPDs and PWDs of the CCS reads that feature a perfect alignment over the windows. We then constructed a kinetic signal pileup using all the IPD or PWD stretches corresponding to the windows. For each CpG, the mean kinetic signal (IPD or PWD) was computed by constructing a pile-up using the kinetic stretches falling within the CpG window and computing the per-column averages. The presence of 5mC created specific IPD (Fig. 1A) and PWD signatures (Fig. 1B).

**Fig. 1.**
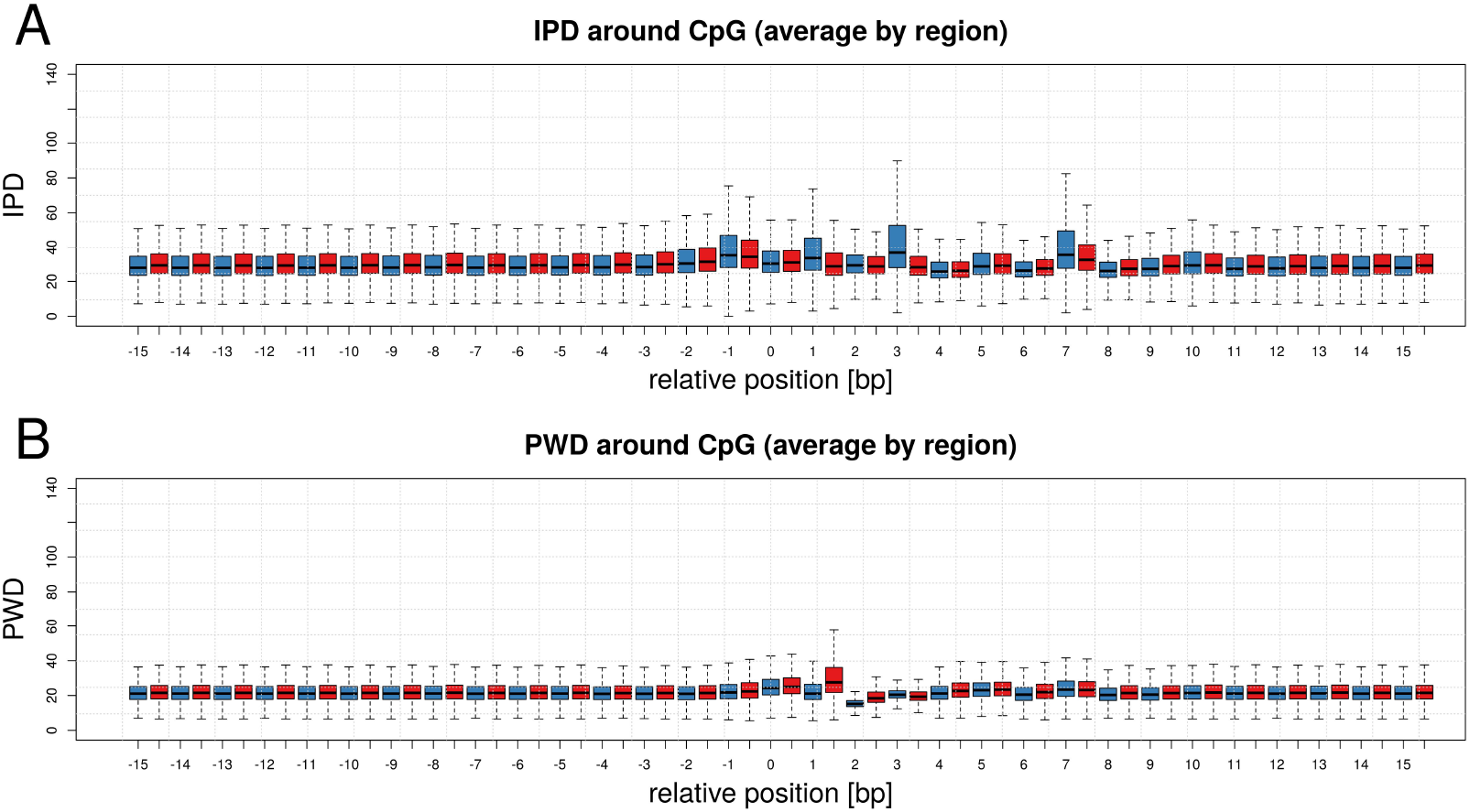
Kinetic signal around CpGs. **A**: Distribution of IPD signal in a 31bp window centered on all CpGs located on chromosome 1 (C is located at position +/-0). The IPD signal was extracted in a per CpG manner. For a given CpG, all CCSs mapping perfectly over the window were considered. The IPD signal was extracted from the positions within the CCS corresponding to the window and the per position averages were computed. This process was repeated for all CpGs on chromosome 1 and the distributions of values is displayed per position. The red and blue distributions indicate the WGA and SssI samples respectively. **B**: Same as A) but with the PDW signal.

We observed an increase of IPD values at position -1, +1, +3 and +7 and a decrease of the PWD signal at positions +1 and +2 relative to the CpGs in the positive sample. The signature was clearer when considering the per-CpG kinetic averages (Fig. 1) rather than simply the CCS kinetics (Fig. S1). The latter corresponds to a reference-free method where it suffices to identify the CpGs within the unmapped CCSs and extract the IPD and PWD kinetics falling within the 31bp windows. We concluded that these differences underline the benefits of mapping the CCSs to study CpGs methylation.

We sought to verify if there was any inter-position dependencies within the window in the positive sample. We measured this by computing the correlation between the kinetic signal seen at any pair of positions within the window (Fig. 2). This revealed strong IPD kinetic dependencies between positions +2 and +3, +6 and +7, which were also important positions for the signature. There were also less strong correlations between many other positions. We observed this between pairs of positions zipping away from one another (diagonals in the matrix). Dependencies were also observed for the PWD kinetics. Finally, the inter-dependencies were stronger when considering the per-CpG averages rather than simply CCS kinetics (Fig. S2), once underlining the advantage of a reference-based approach. We hypothesised that these positions dependencies reflect the interactions between the DNA template and the DNA polymerase or DNA unwinding efficiency fluctuations [31].

**Fig. 2.**
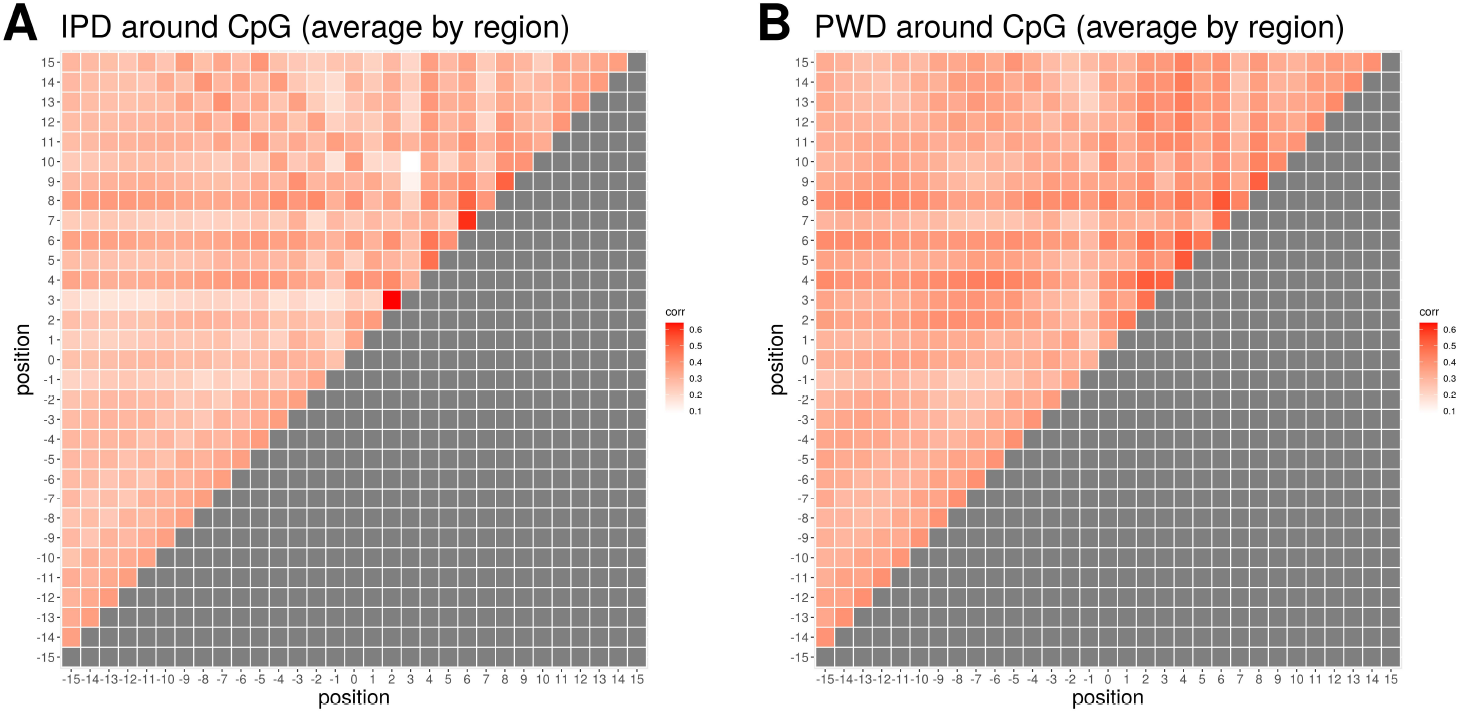
Kinetic signal correlation around CpGs. **A**: Correlation of the IPD signal, averaged per CpG, between all pairwise positions within a 31bp window centered on CpGs (C is located at position +/-0) from chromosome 1, 2 and 3, from the positive sample. For a given CpG, all CCSs mapping perfectly over the CpG window were considered. The IPD signal was extracted from the positions within the CCS corresponding to the window and the per position averages were computed. **B**: Same as A) but with the PDW signal.

### 2.2 Kinetic signal models

Based on the previous observations, we decided to model the observed mean IPD and PWD kinetics of PacBio CCS mapping over a CpG as a sampling from a mixture of two kinetic signal classes: i) methylated and ii) unmethylated CpG signal classes.

Let *R*^+^ be a region of size |*R*^+^|bp on the forward strand and centered on a CpG. *R*^+^ is obtained by considering the position of the C in the CpG and extending it on both sides by *L*bp, resulting in |*R*^+^| = 2*L* + 1bp. This window contains all positions 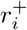 with *i* ∈ (−*L*, −*L* + 1, …, 0, …, *L* + 1, *L*) where 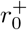 and 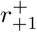 correspond to C and G respectively. The same is applied to the CpG on the reverse strand to obtain *R*^−^. Because |*R*^+^| = |*R*^−^|, in the text below we will refer to the size of the region as |*R*|. *R*^+^ and *R*^−^ do not perfectly overlap, they have a 1bp overhang at each extremity due to the 1bp shift between the C on the + and - strands.

Let *C* = {*c*^1^, *c*^2^, …, *c*^*n*^} be the set of CCSs mapping perfectly - without missmatch nor indel - over *R*^+^ and *R*^−^. Each CCS *c*^*j*^ has a length of |*c*^*j*^| bp.

A CCS is a consensus sequence obtained from the sequencing of both strands of the DNA. Hence, it contains kinetics for both *R*^+^ and *R*^−^. Let *i*^*j*,+^ be defined as the vector of IPD kinetics of *c*^*j*^ corresponding to the forward strand. It is possible to extract a sub-vector 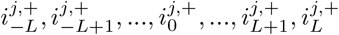 corresponding to the IPD kinetics mapping over *R*^+^. The same can be done for the reverse strand kinetics *i*^*j*,−^, resulting in a sub-vector 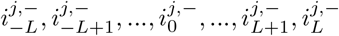 corresponding to the IPD kinetics mapping over *R*^−^.

As previous work has shown [27], two CpGs facing each other on both DNA strands share a consistent methylation state, because of the action of DNMT1. Thus, we decided to combine the information on both strands in order to improve the kinetic information. For this end, we simply decided to link each position 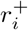 with its counterpart on the other strand 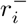. From here onward, CpG will refer to the paired complementary CpGs unless stated otherwise.

This allows us to construct a pileup *W* of IPD kinetics sub-vectors. *W* has 2 * |*C*| rows and |*R*| columns. The left-most column of *W* corresponds to the 5’-most IPD kinetic of each sub-vector and the right-most column to the 3’-most kinetic.

Computing the per column averages of *W* provides us with the strand-aggregated average IPD signal 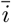 measured for this CpG. The same can be done with the PWD kinetic values to obtain the strand-aggregated average PWD signal 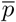

### 2.3 Predicting the presence of 5mC

Let us assume we are provided with four sets of |*R*| distributions 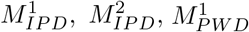 and 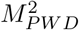 Let 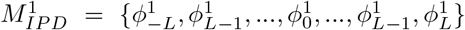 and 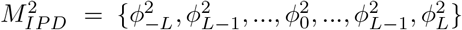 describe the distribution of IPD kinetics expected at every position in a window if the CpG is methylated 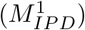 or not 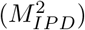 Similarly, 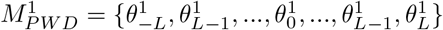and 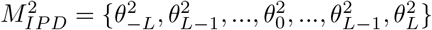 describe the PWD kinetics. This allows to compute the likelihood of IPD or PWD kinetics originating from each of the signal classes as:

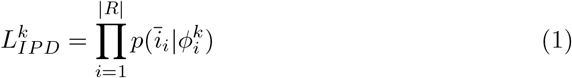

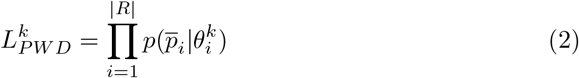

where *k* = 1 for the methylated case and *k* = 2 for the unmethylated case. Because both 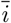 and 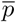 were computed from the same window *R*, we can compute the overall likelihood of observing these IPD and PWD kinetics, given a class model as:

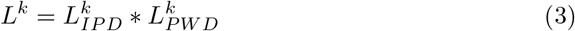

In turn, we can compute the posterior probability of a window containing a methylated CpG:

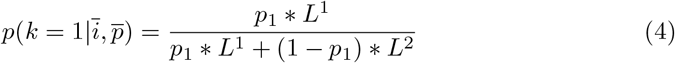

where *p*_1_ is the a-priori probability of this CpG to be methylated. In other word, this is the probability that this window bears a 5mC in its center, given its kinetic signal.

### 2.4 Accounting for pair-wise dependencies

The above model assumes that the kinetic signal at each position is independent from the kinetic signal at any other position in the window. However, as shown in Fig. 2, this assumption does not stand. To account for this, we modified the formulation of kinetic signal class models and turned them into 2-dimension distributions. Each distribution models the kinetic signal at position *x* in the window with respect to the kinetic signal at another position *y* in the window with *x* ≠ *y*.

We designed two types of models: 1) ”di-position” (DP) models accounting only for adjacent position dependencies and 2) ”pair-wise” (PW) kinetic models, accounting for all possible pair-wise dependencies in the window.

DP models account for the set of neighbour position dependencies *D*_*MD*_ = {(−*L*, −*L* + 1), (−*L* + 1, −*L* + 2), …, (*L* − 1, *L*)}, for a total of |*D*_*MD*_| = |*R*| − 1 interactions. The models for the methylated and unmethylated classes are defined as 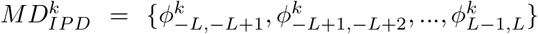 where 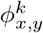 designates the distribution of IPD kinetics, for class *k*, for position *x* with respect to position *y* in the window. Similarly, the models for the PWD kinetics are 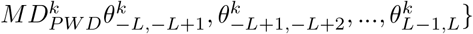.

PW models account for the set of position dependencies *D*_*MP*_ = {(−*L*, −*L* + 1), (−*L*, −*L* + 2), …, (−*L, L*), (−*L* + 1, −*L* + 2), …, (*L* − 1, *L*)}, for a total of 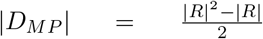 interactions. The models for the methylated and unmethylated classes are defined as the sets 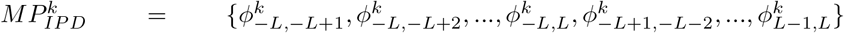 Similarly, the models for the PWD kinetics are 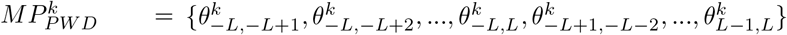.

From this, we can modify equations 1 and 2 as follows:

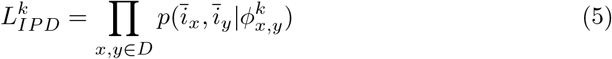

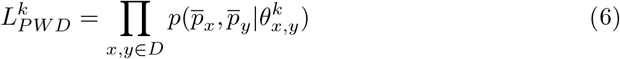

where *D* is *D*_*MD*_ and *D*_*MP*_ for DP and PW models respectively. The overall class likelihood and the probability of CpG methylation are computed using equations 3 and 4.

### 2.5 Sequence effect normalization

Previous studies have shown that the DNA sequence itself, irrespective of DNA modifications, creates fluctuations in the kinetic signal [18, 27]. This is obviously a problem for a classifier. We thus decided to assess the effect of the sequence on the kinetic signal. For this, we computed the average IPD signal, from unmethylated CCSs, in a kmer specific way. Under the assumption that the underlying DNA sequence has no effect on the kinetic signal, we expected the average IPD or PWD measured for each kmer to be somewhat uniform. When displaying the average signal per 7-mer, we immediately noticed considerable variations across position of individual kmers and across kmers (Fig. 3A and Fig. S3A). We then computed the average IPD or PWD per 7-mer from the methylated sample CCSs and divided them by the non-methylated corresponding average values (Fig. 3B and Fig. S3B). IPD and PWD ratios (IPDr and PWDr) displayed a diminished variance across both positions, for a given kmer, and kmers. Additionally, the strongest IPDr / PWDr variation left was strongly correlated with the presence of CpG in the 7-mers. This signal is the variation caused by the presence of 5mC alone, without the effect of the sequence.

**Fig. 3.**
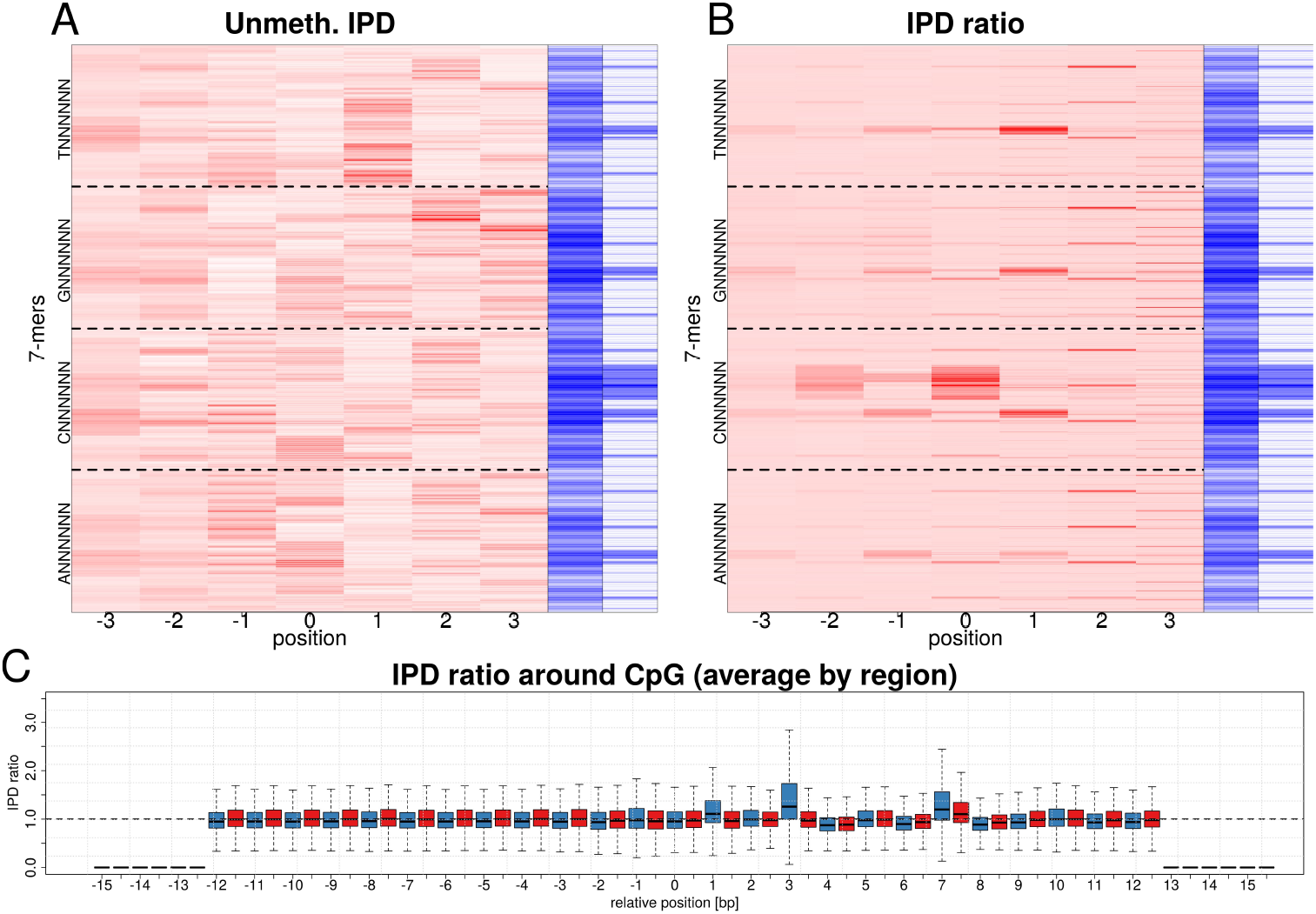
Effect of the sequence on the IPD signal. **A**: Heatmap of the mean IPD signal at each position of all possible 7-mers, from the WGA sample CCSs. The corresponding sequences and IPD signals were extracted and the mean IPD signal was computed for each 7-mer. The 7-mers are sorted in lexicographic order from bottom to top. The dashed lines indicate the limits between A, C, G and T starting 7-mers. The blue bars indicate the 7-mer CG content (center) and CpG content (right) respectively. **B**: Heatmap of the mean IPD ratios (IPDr) at each position of all possible 7-mers. The mean per 7-mer IPD values were computed from the SssI sample CCS as describe in A). Each 7-mer SssI mean IPD was then divided by the corresponding 7-mer WGS average. The ordering, dashed lines and blue bars are the same as in A). **C** Distribution of IPDr signal in a 31bp window centered on all CpGs located on chromosome 1 (C is located at position +/-0). The IPDr signal was extracted in a per CpG manner. For a given CpG, all CCSs mapping perfectly over the CpG window were considered. The IPD signal was extracted from the positions within the CCS corresponding to the window, normalized using a 7-mer backgound model and the per position averages were computed. This process was repeated for all CpG on chromosome 1 and the distributions of values is displayed per position. The first and last six positions in the window have no signal because the normalization process trims the signal, as explained above. The red and blue distributions indicate the WGA and SssI samples respectively.

In order to account for this, we devised a kinetic normalization method (Fig. S4B). For this, we needed to know the expected kinetic signal if there was nothing else but the DNA sequence to dictate kinetic fluctuations.

Let *N* be a map with 4^*K*^ entries corresponding to every DNA sequence *S* of length *K*. Each sequence *S* is mapped to its expected IPD and PWD kinetics *n*_*S,IP D*_ and *n*_*S,P W D*_ respectively of size *K*. Each vector containing the average expected IPD and PWD kinetic at each position of 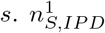 corresponds to the IPD value expected when sequencing the 1st position of 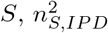 to the 2nd and so one.

To normalize the IPD kinetics, we need to create two independent pileups *W* ^+^ and *W*^−^ with the IPD kinetics mapping over *R*^+^ and *R*^−^ respectively.

Let *S*^+^ and *S*^−^ be the sequences of *R*^+^ and *R*^−^ respectively. There are |*R*|− *K* + 1 IPD sub-vectors of length *K* inside *R*^+^ and *R*^−^ respectively, starting at positions *j* = −*L*, −*L* + 1, …, *L* − *K* + 1 with respect to the CpG. Each of these IPD sub-vector is associated with a corresponding sub-sequence starting at the same position *j*. In turn, *N* contains the expected IPD and PWD values for each given sub-sequence of length *K*. Thus, it is possible to normalize each IPD and PWD sub-vector, in both *R*^+^ and *R*^−^, by dividing them term by term with their expected counters parts.

To normalize a vector of kinetics *Z* of length |*Z*| *> K*, each sub-vector of length *K* is considered independently. The central element of the sub-vector, at position 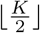, is normalized. Consequently, the edges of *Z* are not normalized and the normalized vector is smaller than the original and has length 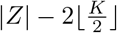.

When we recomputed the kinetic signatures within a 31bp window around CpGs, using a 7-mer normalization, this led to improved 5mC signatures (Fig. 3C and Fig. S3C).

With this information at hand, we decided to train sequence background models with *K* of 7, 9 and 11bp. Low values of *K* are not informative whereas high values of *K* result in less precise average estimates because each sub-sequence is present a limited number of time or even absent from the dataset.

### 2.6 Model training

We decide to model the kinetic signal distributions empirically and used 1-dimension histograms for models without- and 2-dimensional histograms for models with inter-dependencies. Training a methylated kinetic signal class model requires a set of CCSs for which we a certain of the methylation state. Similarly, training an unmethylated kinetic signal class model requires CCSs in which we have the certainty that there is no DNA modification. We trained one model from the HG002 SssI sample to estimate the methylated CpG kinetic signal and one model from the HG002 WGA sample to estimate the unmethylated CpG kinetic signal. The training process requires mapping the CCSs against a genome of reference and to list all CpGs within this genome.

The following steps are applied to each CpG of the genome independently (Fig. S5). A window of size |*R*| bp (|*R*| is odd), around a CpG, centered on the C, can be built. The sub-vector of IPD kinetics mapping inside the window are extracted for both strands independently. The kinetic pileup *W* can be build from these kinetic sub-vectors. The per-column average yields the average per-position IPD 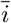 for this window.Each value 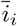 is fed to 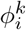 inside 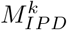 The exact same procedure is followed for 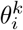 inside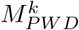, with the PWD kinetic.

Training DP and PW models follows the same logic (Fig. S6 and Fig. S7) until the creation of the per-position average IPD and PWD kinetic signals 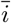 and 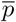 Each pair of values at positions *x, y* ∈ *D*_*DP*_ for DP models and *x, y* ∈ *D*_*P W*_ for PW models is considered and fed to the corresponding 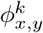 and 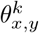 histograms respectively.

We also trained each type of model using a sequencing normalizations of the IPD and PWD kinetics. The procedure is exactly the same as described above with the following exceptions. First, the window around the CpG is extended to 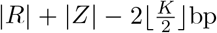 in order to get exactly |*R*| normalized stretches (see above). Second, the IPD and PWD kinetics are normalized before being fed to the histograms.

We decided to train models of 17bp length (+/-8bp around the C of the CpG). We deemed this window size sufficient to capture most of the kinetic signature information since the most important distribution differences were observed between positions -1 and +7 (Fig. 1 and Fig. 2). Models were trained by pairs: one from the WGA sample to model the unmethylated kinetic signal and one from the SssI sample to model the methylated kinetic signal.

We trained a total of 12 different pairs of model, to cover all possible cases : 1) position independent models (K=7,9 and 11bp) with and without sequence normalization, 2) PW models (K=7,9 and 11bp) with and without sequence normalization, and 3) DP models (K=7,9 and 11bp) with and without sequence normalization.

### 2.7 CpG methylation prediction power

We then assessed the prediction power of these models using the Telomere-To-Telomere Consortium (T2T) generated data. This dataset contains SequelII data with a genome-wide average 42X coverage [32, 33] as well as a methylation track constructed from ONT sequencing data [34]. Recent work has demonstrated that ONT data can be used to confidently predict CpG methylation. Thus, we decided to use this track as our truth set. As genome reference, we decided to use the CHM13v1.0 genome assembly by T2T [35]. To date, this assembly is the most complete human reference, with the most comprehensive delineation of complex/repeated regions.

We listed all the CpGs present in the CHM13v1.0 genome. For each CpG, we extracted the kinetics falling within a 17bp window centered on the CpG from every CCS mapping perfectly in the window. We then computed the window average IPD and PWD values and fed them to the predictor models.

For comparison, we also computed predictions using PacBio’s pb-CpG-tools. Any CpG with a methylation probability *>* 0.5 was considered methylated, conversely any CpG with a methylation probability *<* 0.5 was considered unmethylated. We then compared the predictions of all models against the T2T methylation track (Fig.4). PacBio’s pb-CpG-tools achieved an impressive Area Under the Curve (AUC) of 0.97 with a high sensitivity (also called recall). Our models showed a wide variety of performances. The kinetic normalization clearly improved the performances, bringing an AUC of 0.83 for simplest model to an AUC of 0.93 when adding an 11bp background sequence normalization. Accounting for inter-position dependencies improved the specificity of the models, with all our models achieving a higher sensitivity than pb-CpG-tools. The best model on our side (11bp normalization and di-position modelling) achieved an AUC of 0.93, lower than pb-CpG-tools, yet showing a superior specificity and precision with 0.94 and 0.93, respectively. This indicate that PacBio’s predictor is better at finding methylated CpG but at the expense of wrongly calling methylated CpGs.

**Fig. 4.**
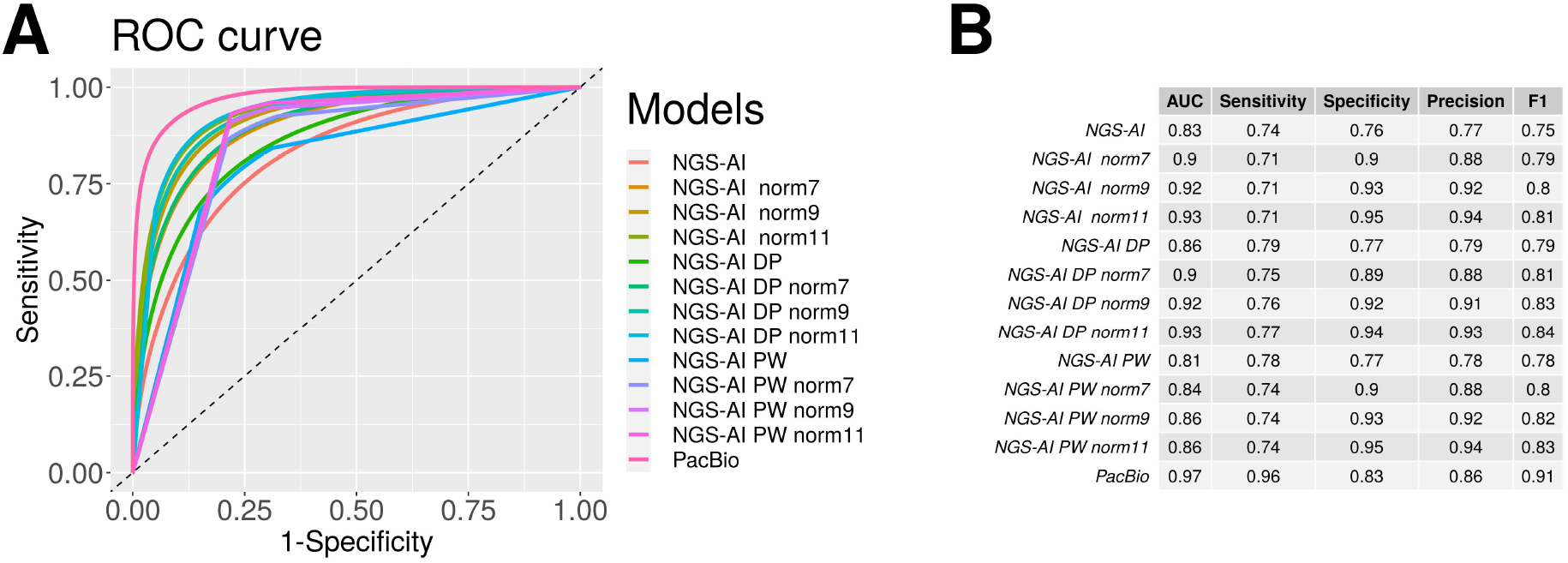
Classification performances. **A**: the ROC curves for each model. The classification performances were assessed using T2T methylation track as truth set. The models labelled as “norm-7”, “norm-9” and “norm-11” used a sequence normalization background with 7-mers, 9-mers and 11-mers, respectively. Models labelled as “DP” and “PW” accounts for direct neighbour dependencies and all pairwise dependencies within the window, respectively. **B**: The detailed performance statistics per model.

Finally, we assessed the impact of coverage on the prediction performances of the best models (pb-CpG-tools and di-position models with 11bp normalisation). For this, we down-sampled mapped CCSs down to a coverage of 20X, 10X, 5X, 2X and 1X and computed the predictions (Fig. 5). Overall, the same trend was observed with decreasing coverage. pb-CpG-tools model showed better AUC and specificity but had a lower sensitivity and precision. Our model started showing a better AUC with extremely sparse sequencing coverage of 2X and 1X.

**Fig. 5.**
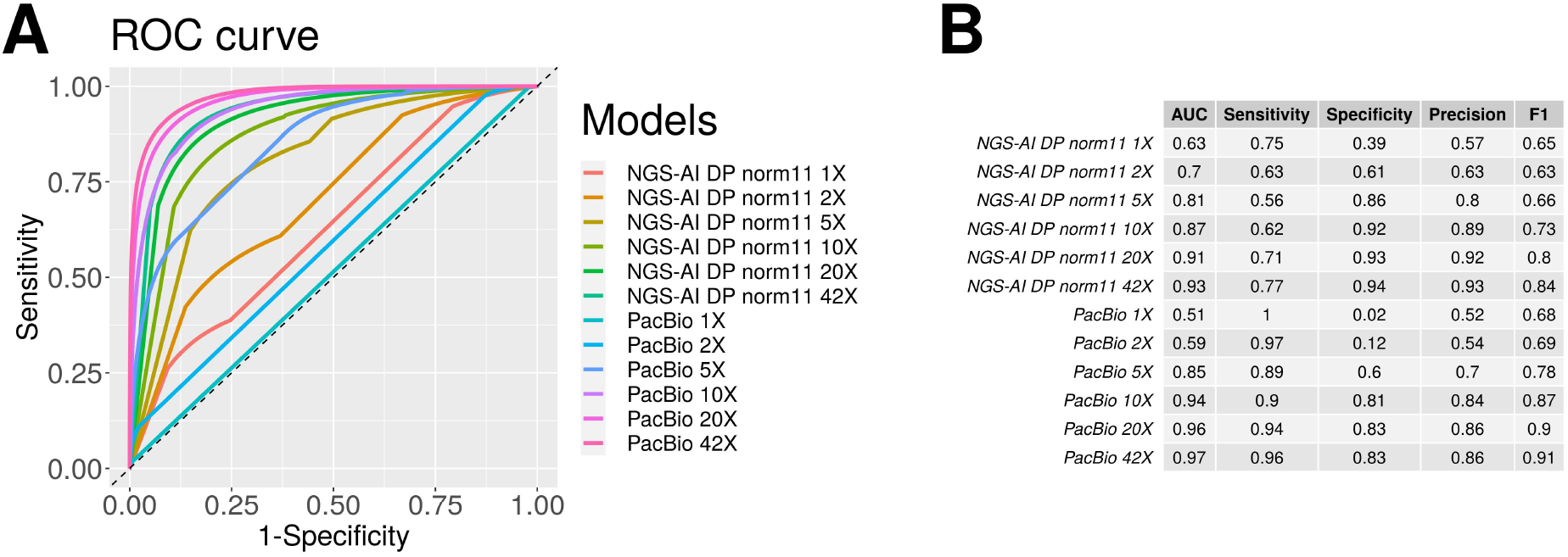
Effect of the coverage on the classification performances. **A**: classification performances achieved by the best models for different coverages. The classification performances were assessed using T2T methylation track as truth set. The whole dataset (42X) was down-sampled in order to obtain global average coverage of 1X, 2X, 10X, and 20X. The models labelled as “norm-7”, “norm-9” and “norm-11” used a sequence normalization background of 7-mers, 9-mers and 11-mers respectively. Models labelled as “DP” accounts for direct neighbour dependencies within the window respectively. **B**: The detailed performance statistics per model.

## 3 Discussion

We showed that 5mC creates specific IPD and PWD kinetic signatures around CpGs (Fig. 1). We also showed that specific positions around the CpG resonate together in the kinetic signatures (Fig. 2 and that the sequence background has a strong influence on the kinetics (Fig. 3A and Fig. S3A). From this, we have designed a framework to model this signal and train the models from the data. Finally, we designed several types of signal models : normal models, DP models accounting for adjacent position inter-dependencies and PW models accounting for any two position inter-dependencies.

Atop of this, we designed a way to normalize the signal for each model to account for the effect of the sequence on the kinetics.

Overall, our models showed similarly good performance as other available methods. PacBio’s primerose tool showed a better ability to predict the presence of 5mC at CpG, at the expense of falsely calling 5mC when there is none. However, our di-position normalised model showed a better propensity to distinguish unmethylated CpGs and made less mistakes than PacBio’s tool when predicting 5mC.

There are a few aspects which hinder potentially at the moment a superior performance. First, in the training data, the 5mC is not really removed from the DNA in the WGA sample, but is rather diluted out. Consequently, a small fraction of the allegedly unmethylated CpG contain still 5mC, most likely on a single strand. Similarly, the SssI mediated methylation efficient is only ∼95% [36]. Some of these CpGs are thus left unmethylated in the positive control sample. Consequently, both the positive and negative datasets contain residual CpGs in the opposite state. This is expected to introduce a bias in the model, that latter impacts the classification performances. and can lead to discrepancies compared with WGBS or ONT inferred methylation. Second, the presented approach is not allele-aware. Therefore it does not distinguish different alleles and likely miss-classifies CpG falling in differently methylated alleles by providing an intermediate score. This can potentially be mitigated by phasing and separating the alignments but has not been assessed in this work.

Furthermore, we think there is still room to improve the predictor performances through the optimization of the model and parameters. For instance, we have not tested the effect of the window size. The same is true for the prior probability of a CpG to be methylated. In this work, we used a neutral choice of 0.5. However there may be situations in which adapting it can turn useful, for instance in case of deficient methylation deposition pathways, or on a specie-specific basis.

These algorithms are general enough to allow a wide variety of usages. First, we did not introduce any specie specific parameters and PAPET could therefore be used for non-human data as well. Second, this framework and tools could be used to train a predictor against any other epigenetics marks, such as N6-methyladenosine if comparable data are available.

## 4 Conclusions

With this work, we showed that 5mC creates specific IPD and PWD kinetic signatures around CpGs (Fig. 1). We also highlighted that certain specific positions around the CpG resonate together in the kinetic signatures (Fig. 2). Finally, we also demonstrated the effect of the DNA sequence content on the kinetics (Fig. 3A and Fig. S3A).

We propose a framework to model the PacBio kinetic signal and have designed a kinetic normalization to account for the DNA sequence effect. From this, we developed and trained different families of predictive models, with- or without position inter-dependencies, and demonstrated that they perform as good as their best in class in terms of classification performances. Finally, the data, softwares and prediction models have been made readily available for the research community.

## 5 Availability

### 5.1 Data

The following table lists all the datasets used for this project as well as their accession numbers on the SRA database.

| Origin | Accession | Individual | Treatment |
| --- | --- | --- | --- |
| PacBio | SRR24523544 | HG002 | WGA |
| PacBio | SRR24523543 | HG002 | WGA |
| PacBio | SRR24523540 | HG002 | WGA |
| PacBio | SRR24523539 | HG002 | WGA |
| PacBio | SRR24523533 | HG002 | SssI |
| PacBio | SRR24523534 | HG002 | SssI |
| PacBio | SRR24523535 | HG002 | SssI |
| PacBio | SRR24523536 | HG002 | SssI |
| PacBio | SRR24523537 | HG002 | SssI |
| PacBio | SRR24523538 | HG002 | SssI |
| PacBio | SRR24523541 | HG002 | SssI |
| PacBio | SRR24523542 | HG002 | SssI |
| T2T | SRR11292120 | CHM13 | None |
| T2T | SRR11292121 | CHM13 | None |
| T2T | SRR11292123 | CHM13 | None |

### 5.2 Code and software

PAPET source code is available at https://github.com/ngs-ai-org/papet. A docker image with PAPET software suite is available at https://hub.docker.com/r/ngsai/papet.

### 5.3 Models

The 5mC prediction models trained are available at https://github.com/ngs-ai-org/papet-models.

## 6 Abbreviations

CpG: cytosine followed by a guanine
5mC: 5-methylcytosine
CGI: CpG island
DNMT: DNA-methyl-transferase
5hmC: 5-hydroxymethylcytosine
5fC: 5-formylcytosine
TSS: transcription start site
H3K4me3: H3 lysine 4 tri-methylation
H3K36me2: H3 lysine 36 di-methylation
RNAPII: RNA polymerase II
H3K27me3: H3 lysine 27 tri-methylation
WGBS: whole genome bisulfite sequencing
WGS: whole genome sequencing
SMRT-seq: single molecule real-time sequencing
ONT: Oxford Nanopore Technologies
PacBio: Pacific Biosciences
PWD: pulse width
IPD: interpuse duration
IPDr: IPD ratio
PWDr: PWD ratio
WGA: whole genome amplified
HMM: hidden Markov model
RNN: recurrent neural network
CNN: convolutional neural network
LSTM: long short-term memory
LDA: linear discriminant analysis
CCS: circular consensus sequence
BiGRU: bidirectional gated recurrent unit
PAPET: PacBio prediction epigenetics toolkit
HiFi: High fidelity

## 7 Material and Methods

### 7.1 Data

For this study, we used two publicly available datasets.

The first dataset was created by PacBio. It originates from the HG002/NA24385 individual. It is composed of 2 samples, referred to as WGA and SssI samples. These samples contain 4 and 8 SMRT-cell respectively. The WGA sample contains genomic DNA that was subjected to WGA prior being sequenced on a PacBio SequelII sequencer. The Sssl sample is derived from the WGA sample by subjecting it to a SssI enzyme treatment prior to sequencing on a PacBio SequelII sequencer.

The second dataset was created as part of the T2T Consortium work. It originates from the human CHM13 cell line. The genomic DNA was extracted from the cells and sequenced with a PacBio SequelII sequencer. It is composed of 3 HiFi (high fidelity) SMRT-cells. A corresponding CpG methylation track derived from ONT sequencing data for the CHM13v1.0 assembly was downloaded from here. The bigWig file was then converted to bedgraph with bigWigToWig using:

### 7.2 Extraction of PacBio CCS kinetic data for analysis

The negative and positive sample CCSs were mapped against hg38p8 genome assembly with `pbmm2` [37] using:

~~~
pbmm2 align --preset CCS \
--sample <sample-name> \
-j 30 --log-level INFO \
<file.in.bam> \
<file.mmi> \
<file.out.bam>
~~~

where `<sample-name>` is the sample label, `<file.in.bam>` is the path to the input bam file, `<file.mmi>` is the path to the genome index file and `<file.out.bam>` is the path to the output bam file.

The coordinates of the CpGs in the hg38p8 genome were found with `getCpG.py`, a program of ours, using:

~~~
python3.6 getCpG.py --fasta hg38_p8.fa | \
sort -k1,1V -k2,2n -k 3,3 \
    > hg38_p8_cpg.bed
~~~

Then kinetic signal of each CCS, around the CpG sites, were extracted with `papet kinetics`, using:

~~~
papet kinetics
--bed hg38_p8_cpg.bed \
--bam <file_list> \
--winSize 31 > kinetics.tsv
~~~

where `<file_list>` is a coma-separated list of bam files containing the CCS from which the kinetic will be extracted. `papet kinetics` was run once with all the WGA sample bam files and once with all the Sssl sample bam files.

The same procedure was used to extract normalized kinetic signal:

~~~
papet kinetics \
--bed hg38_p8\_cpg.bed \
--bam <file_list> \
--winSize 31 \
--model <file_model_bckg> > kinetics_normalized.tsv
~~~

where `<file_model_bckg>` contains the background sequence model to use to normalize the kinetics. Three different models were used : 7bp 9bp and 11bp.

### 7.3 Background sequence training

The background sequence models were trained using the WGA sample data with `papet model-sequence`, using:

~~~
papet model-sequence \
--bam <file_list> \
--kmer <k> \
--out <file_model_bckg>
~~~

where `<file_list>` is a coma-separated list of bam files containing non-methylated CCS, `<k>` is the kmer size to use and `<file_model_bckg>` is the path where the sequence background model will be written.

### 7.4 Kinetic model training

To train the kinetic signal models, we used the Pacific Bioscience dataset. The models were always created by pairs. One model from the WGA sample data describing the expected unmethylated CpG signal without and one model from the Sssl sample data describing the expected methylated CpG signal.

The simplest models, accounting for neither inter-position dependencies nor the DNA sequence effect, were trained with `papet model-kinetic`, using:

~~~
papet model-kinetic \
raw \
--bed hg38_p8_cpg.bed \
--bam <file_list> \
--size 17 \
--nbin 952 \
--xmin 0 \
--xmax 952 \
--pseudocount 1 \
--out <file_model>
~~~

where `<file_list>` is a coma separated list of all bam file paths from either the negative or positive sample, `--nbin` is the number of bins inside the histograms, `--xmin` is the lower boundary of the lower bin in the histograms, `--xmax` is the upper boundary of the upper bin in the histograms, `--pc` is a pseudocount value added once to each histogram bin to avoid 0 count bins and `<file_model>` is the path where the kinetic model will be written. `--nbin`, `--xmin` and `--xmax` were set to 952, 0 and 952 because raw IPD and PWD values can take any integer value from 0 to 952.

Normalized kinetic models, accounting for the DNA sequence effect only, were trained with `model-kinetic`, using:

~~~
papet model-kinetic \
raw-norm \
--bed hg38_p8_cpg.bed \
--bam <file_list> \
--background <file_model_bckg> \
--size 17 \
--nbin 952 \
--xmin 0 \
--xmax 5 \
--pseudocount 1 \
--out <file_model>
~~~

where `<file_model_bckg>` is the path to a sequence background model created with `model-sequence`. We used three different sequence models : 7bp, 9bp and 11bp. The values of `--xmin` and `--xmax` were set to 0 and 5 because normalized IPD and PWD values cannot be smaller than 0 and were mostly smaller than 5. In addition, two bins per histogram dimension were automatically added to cover the inverals ]−∞,0] and ]5,∞[. Finally, `--nbin` was set to 952 in order to obtain the same level of granularity as with the simplest model.

PD kinetic models, accounting for neighbouring position inter-dependencies only, were trained with `model-kinetic`, using:

~~~
model-kinetic \
diposition \
--bed hg38_p8_cpg.bed \
--bam <file_list> \
--size 17 \
--nbin 100 \
--xmin 0 \
--xmax 100 \
--pseudocount 1 \
--out <file_model>
~~~

The value of `--xmin`, `--xmax --nbin` were set to 0, 100 and 100. Since these histograms are 2-dimensional, they contain 100 * *2 bins. Since both the IPD and the PWD are smaller than 100, we created integer bins from 0 to 100. In addition, two bins per dimension were automatically added to account for the values passed the min and max of the histograms.

Normalized PD kinetic models, accounting for neighbouring position inter-dependencies and DNA sequence effect, were trained with `model-kinetic`, using:

~~~
papet model-kinetic \
diposition-norm \
--bed hg38_p8_cpg.bed \
--background <file_model_bckg> \
--bam <file_list> \
--size 17 \
--nbin 100 \
--xmin 0 \
--xmax 5 \
--pseudocount 1 \
--out <file_model>
~~~

where `<file_model_bckg>` is the path to a sequence background model created with `model-sequence`. We used three different models: 7bp, 9bp and 11bp. The values of `--xmin` and `--xmax` were set to 0 and 5 because normalized IPD and PWD values cannot be smaller than 0 and were mostly smaller than 5. In addition, two bins per histogram dimension were automatically added to cover the inverals ]−∞,0] and ]5,∞[. Finally, `--nbin` was set to 100. Indeed these histograms are 2-dimensional. They contain 100 * *2 bins, plus two bins per dimension to account for the values passed the min and max of the histograms.

PW kinetic models, accounting for all position inter-dependencies only, were trained with `model-kinetic`, using:

~~~
papet model-kinetic \
pairwise \
--bed hg38_p8_cpg.bed \
--bam <file_list> \
--size 17 \
--nbin 100 \
--xmin 0 \
--xmax 100 \
--pseudocount 1 \
--out <file_model>
~~~

The value of `--xmin`, `--xmax --nbin` were set to 0, 100 and 100. Since these histograms are 2-dimensional, they contain 100 * *2 bins. Since both the IPD and the PWD are smaller than 100, we created integer bins from 0 to 100. In addition, two bins per dimension were automatically added to account for the values passed the min and max of the histograms.

Normalized PW kinetic models, accounting for all position inter-dependencies and DNA sequence effect, were trained with `model-kinetic`, using:

~~~
papet model-kinetic \
pairwise-norm \
--bed hg38_p8_cpg.bed \
--background <file_model_bckg> \
--bam <file_list> \
--size 17 \
--nbin 100 \
--xmin 0 \
--xmax 5 \
--pseudocount 1 \
--out <file_model>
~~~

where `<file_model_bckg>` is the path to a sequence background model created with `model-sequence`. We used three different models : 7bp, 9bp and 11bp. The values of `--xmin` and `--xmax` were set to 0 and 5 because normalized IPD and PWD values cannot be smaller than 0 and were mostly smaller than 5. In addition, two bins per histogram dimension were automatically added to cover the inverals ]−∞,0] and ]5,∞[. Finally, `--nbin` was set to 100. These histograms are 2-dimensional. They contain 100 * *2 bins, plus two bins per dimension to account for the values passed the min and max of the histograms.

### 7.5 CpG methylation predictions

For this part, we used the CHM13 SequelII WGS generated by the T2T consortium. We downloaded the subreads corresponding to entries SRR11292120 (m 64062 190806 063919), SRR11292121 (m 64062 190803 042216) and SRR11292123 (m 64062 190804 172951) on the SRA and also available on T2T github page.

Fist, we computed the CCSs from the subreads with PacBio’s `ccs` using:

~~~
ccs -j 4 \
   --all \
   --hifi-kinetics \
   --report-file ccs_report.txt \
   --report-json ccs_report.json \
   --metrics-json ccs_metric.json\
   subreads.bam \
   ccs.bam\
~~~

where `<subreads.bam>` is the path to the file containing the subreads and `<ccs.bam>` is the path to the file that will contain the CCSs. This step was performed for each subread file independently.

Then, the CCSs were mapped against CHM13 genome assembly v1.0 released by T2T with PacBio’s pbmm2 using:

~~~
pbmm2 \
align \
--preset CCS \
--sample t2t-1 \
-j 30 \
--log-level INFO \
ccs.bam \ chm13.draft_v1.0.mmi \
ccs.chm13v1_0.bam \
~~~

where `<ccs.bam>` is the path to the file containing the CCSs, `<chm13.draft_v1.0.mmi>` is the path to the indexed genome and `<ccs.chm13v1_0.bam>` the path to the file that will contain the mapped CCSs. This step was performed for each CCS file independently. Then all mapped CCSs were pooled together with samtools [38] using:

~~~
samtools merge all.bam \
m64062_190803_042216.ccs.chm13v1_0.bam \
m64062_190804_172951.ccs.chm13v1_0.bam \
m64062_190806_063919.ccs.chm13v1_0.bam && \
samtools sort -o all.sorted.bam
~~~

The coordinates of the CpGs in the hg38p8 genome were found with our program `getCpG.py`, using:

~~~
python3.6 getCpG.py --fasta chm13.draft_v1.0.fasta | \
sort -k1,1V -k2,2n -k 3,3n \
  > chm13.draft_v1.0.cpg.bed
~~~

where `<chm13.draft_v1.0.cpg.bed>` is the file containing the CHM13 v1.0 genome assembly chromosome sequences and `<chm13.draft_v1.0.cpg.bed>` the file containing the genomic coordinates of the CpGs found in the genome.

The predictions using our models were computed with `papet predict`, using:

~~~
papet predict \
--bam all.sorted.bam \
--bed chm13.draft_v1.0.cpg.bed \
--modelMeth <file_model_meth> \
--modelUnmeth <file_model_unmeth> \
--prob 0.5 \
> <predictions.bed>\
~~~

where `<file_model_meth>` and `<file_model_unmeth>` are two files containing models for methylated and unmethylated kinetic signals respectively and `<predictions.bed>` is the file containing the prediction results. Predictions were run using the simplest, the DP, the DP normalized, the PW and the PW normalized models independently.

Predictions using PacBio’s `pb-CpG-tools` were computed using:

~~~
primrose \
--keep-kinetics \
--num-threads 30 \
all.sorted.bam \
all.sorted.primrose.bam && \
samtools index \
-@ 30 \
all.sorted.primrose.bam && \
python /pb-CpG-tools/aligned_bam_to_cpg_scores.py \
-b all.sorted.primrose.bam \
-f chm13.draft_v1.0.fasta \
-o <label> \
--pileup_mode model \
--model_dir /pb-CpG-tools/pileup_calling_model \
--modsites reference
~~~

where `<ccs.bam>` is the path to the file containing all CCSs, `<all.sorted.primrose.bam>` is the file containing the per read methylation predictions computed by `primrose`, `<chm13.draft_v1.0.fasta>` is the file containing CHM13 genome assembly v1.0 as released by T2T. We also tried to run CpG methylation predictions using `ccsmeth` [30]. Unfortunately, we were not successful in running this software.

### 7.6 CCS sub-sampling

To achieve lower coverages, CCS sub-sampling was performed using:

~~~
sambamba view \
-f bam \
-h \
-s <ratio> \
--subsampling-seed <seed> \
all.sorted.primrose.bam > all.sorted.sub.bam \
~~~

where `<ratio>` is a ratio corresponding to the final desired coverage. The 3 pooled SMRT-cells accounted for a total of 42x genome-wide average coverage. In order to sample down to 20x, 10x, 5x, 2x and 1x we used 20/42, 10/42, 5/42, 2/42 and 1/42 respectively. `<seed>` is a seed to set random processes for reproducibility purposes.

### 7.7 Predictor performances

All prediction files were merged into one table containing, for each CpG, all the predictions computed. If a predictor did not produce a prediction for a CpG while at least one other did, the value was set to NA for the missing predictor. For this, we used:

~~~
 python3.6 bedToTable.py <files_pred> > tmp.bed &&
sort -k1,1 -k2,2n -k3,3n tmp.bed > tmp.sorted.bed &&
mv tmp.sorted.bed tmp.bed &&
bedtools map \
 -a tmp.bed \
 -b methyFreq.bedgraph \
 -c 4 -o mean > tmp.tmp.bed &&
sed -E ‘s/\.$/NA/’ tmp.tmp.bed > tmp.bed &&
rm tmp.tmp.bed &&
bedtools closest \
 -D ref -io -a tmp.bed -b tmp.bed \
 cut -f ${echo {1..103} | sed ‘s/ /,/g’},207\
 > chm13.draft_v1.0_cpg_5mC.bed
~~~

where `bedToTable.py` is a program of ours, `<files_pred>` is a coma-separated list of all prediction files created by `papet predict` and `pb-CpG-tools` and `<methyFreq.bedgraph>` is the methylation track released by T2T for CHM13 v1.0. T2T methylation track was considered as the truth set.

The tables were the imported in R. Any CpG for which no truth set methylation value was available were discarded. Missing predictions were set to 0.5, a value that provides no information regarding the methylations status. Finally, CpGs were separated into methylated and unmethylated based on the truth set methylation probability (CpG was considered methylated with *p >* 0.5 and unmethylated with *p <* 0.5). The classifier performances were then measured with the `precrec` R library [39] using:

~~~
pred = mmdata(scores=data, labels=labels)
auc = evalmod(pred)
~~~

where `data` is a dataframe of methylation prediction probabilities with the CpGs on the rows and the predictors on the columns. Labels is a vector of boolean indicating whether a CpG is considered methylated or not according to the truth set.

## 9 Author contributions statement

R.G. designed and implemented the algorithms. R.G. and E.S ran the benchmark and analysed the results. R.G. wrote the manuscript. R.G., I.X. and E.S reviewed the manuscript.

## 10 Acknowledgments

The authors thank Richard Hall and the Pacific Bioscience bioinformatics team for sharing their WGA/Sssl training dataset. Also, the authors express their gratitude to the first author’s mom for sharing the best recipe of papet ever.

## 11 Supplemental Figures

**Fig. S1.**
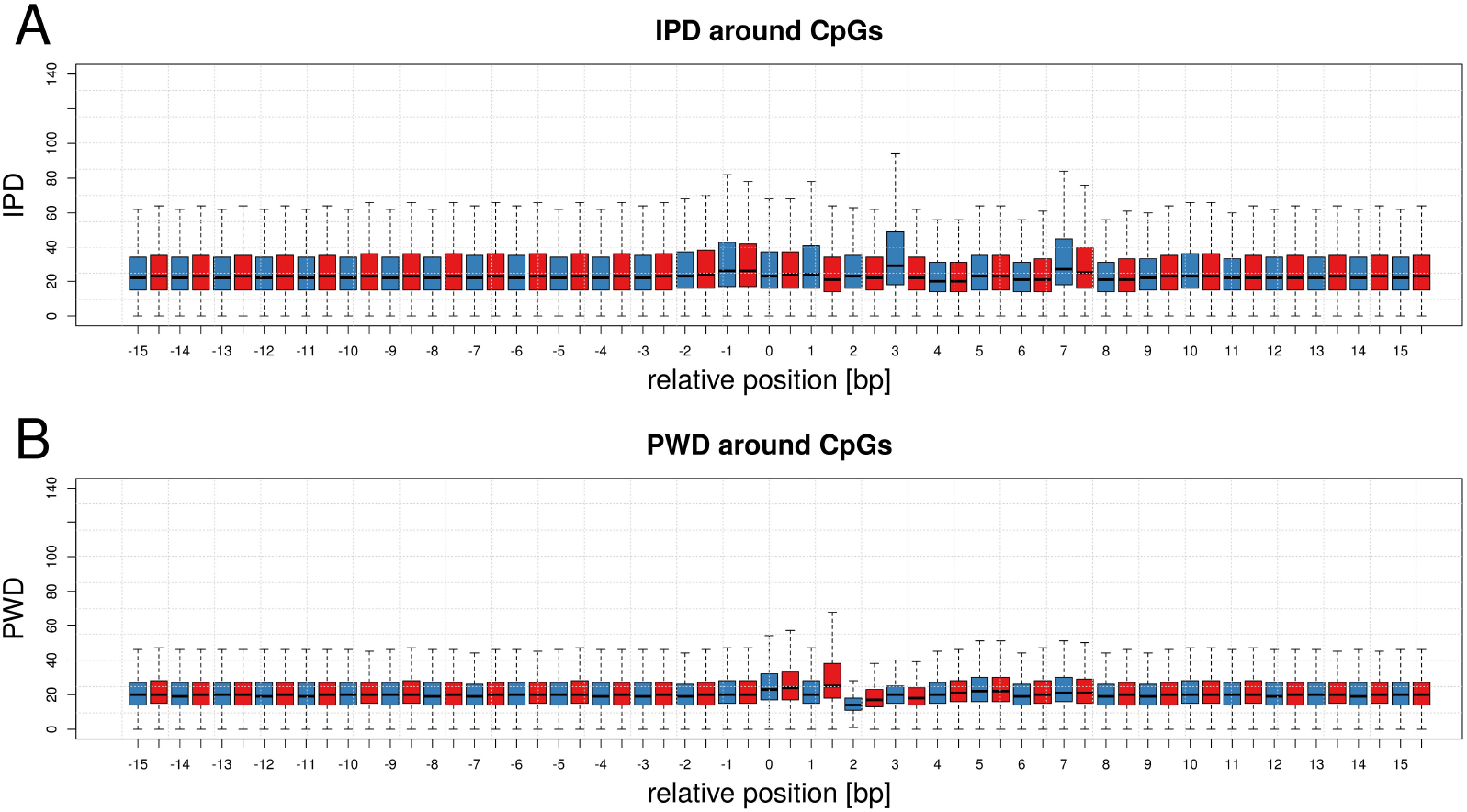
Kinetic signal around CpGs. **A** distribution of IPD signal in a 31bp window centered on all CpGs located on chromosome 1 (C is located at position +/-0). The IPD signal was extracted from the positions within the CCS corresponding to the window. The red distributions indicate CCSs without 5mC, blue distributions CCSs with 5mC. **B**: Same as A but with the PDW signal.

**Fig. S2.**
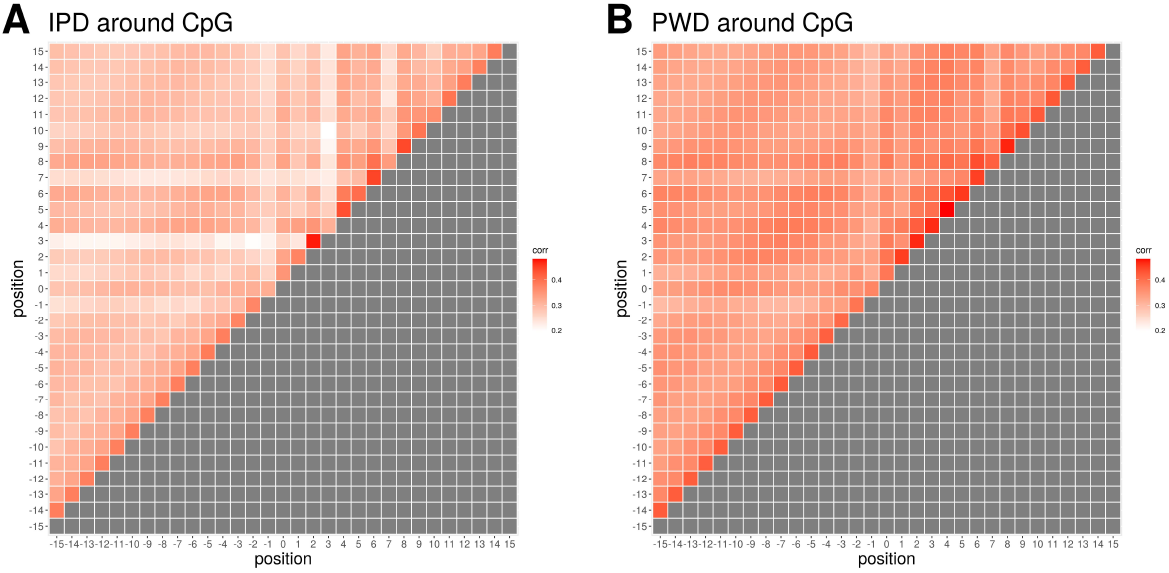
Kinetic signal correlation around CpGs. **A**: Correlation of the IPD signal between all pairwise positions within a 31bp window centered on CpGs (C is located at position +/-0) from chromosome 1, 2 and 3. The IPD signal was extracted from the positions within the CCS corresponding to the window and the per position averages were computed. **B**: Same as A but with the PDW signal.

**Fig. S3.**
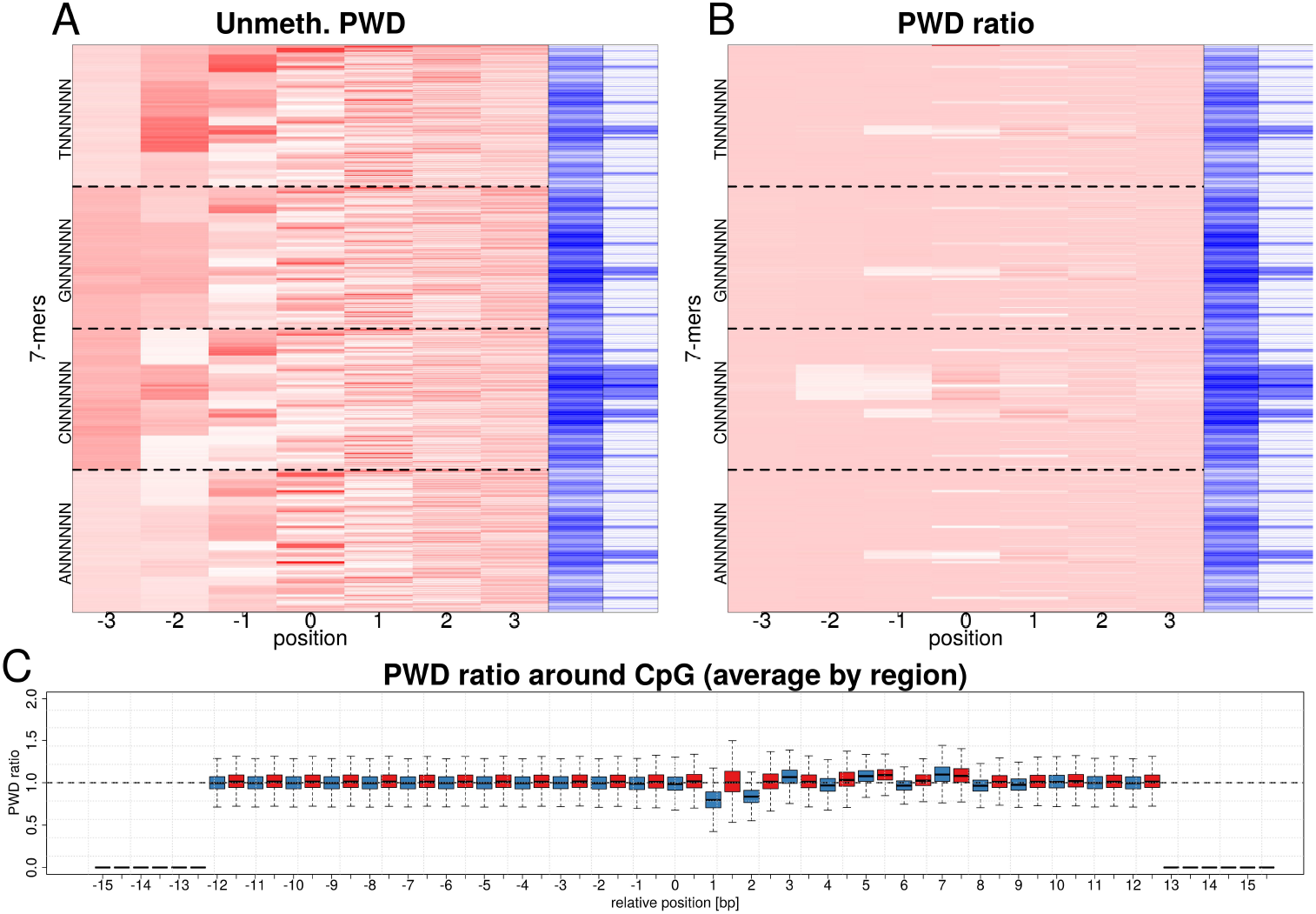
Effect of the sequence on the PWD signal. **A**: Heatmap of the mean PWD signal at each position of all possible 7-mers, from the WGA sample CCSs. The corresponding sequences and PWD signals were extracted and the mean PWD signal was computed for each 7-mer. The 7-mers are sorted in lexicographic order from bottom to top. The dashed lines indicate the limits between A, C, G and T starting 7-mers. The blue bars indicate the 7-mer CG content (center) and CpG content (right) respectively. **B**: Heatmap of the mean PWD ratios (PWDr) at each position of all possible 7-mers. The mean per 7-mer PWD values were computed from the SssI sample CCS as describe in A). Each 7-mer SssI mean PWD was then divided by the corresponding 7-mer WGS average. The ordering, dashed lines and blue bars are the same as in A). **C** Distribution of PWDr signal in a 31bp window centered on all CpGs located on chromosome 1 (C is located at position +/-0). The PWDr signal was extracted in a per CpG manner. For a given CpG, all CCSs mapping perfectly over the CpG window were considered. The PWD signal was extracted from the positions within the CCS corresponding to the window, normalized using a 7-mer backgound model and the per position averages were computed. This process was repeated for all CpG on chromosome 1 and the distributions of values is displayed per position. The first and last six positions in the window have no signal because the normalization process trims the signal, as explained above. The red and blue distributions indicate the WGA and SssI samples respectively.

**Fig. S4.**
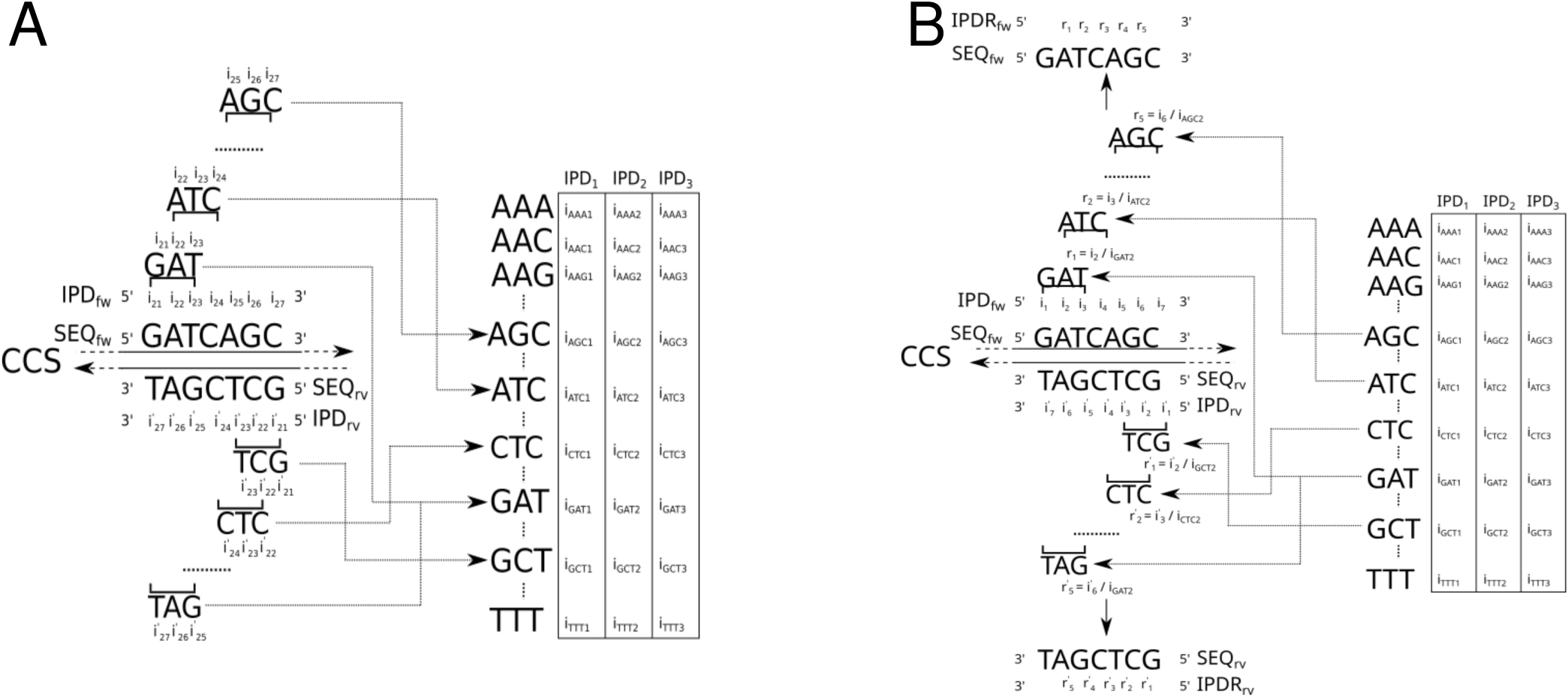
Sequence background model. **A**: A map contains all kmer sequences as keys and the expected per kmer position IPD (or PWD) values without any DNA modification is stored as value. The sequence background model is trained from each kmer in a WGA dataset and computing the per kmer IPD (or PWD) average signal. **B**: A stretch of IPD (or PWD) kinetics can be normalized using the map. The stretch is broken in smaller sub-slices of length *K*. The central position of each sub-slice in the kinetic signal is divided by the central expected IPD (or PWD) value expected for this kmer without DNA modification. This procedure is repeated for every possible sub-slice.

**Fig. S5.**
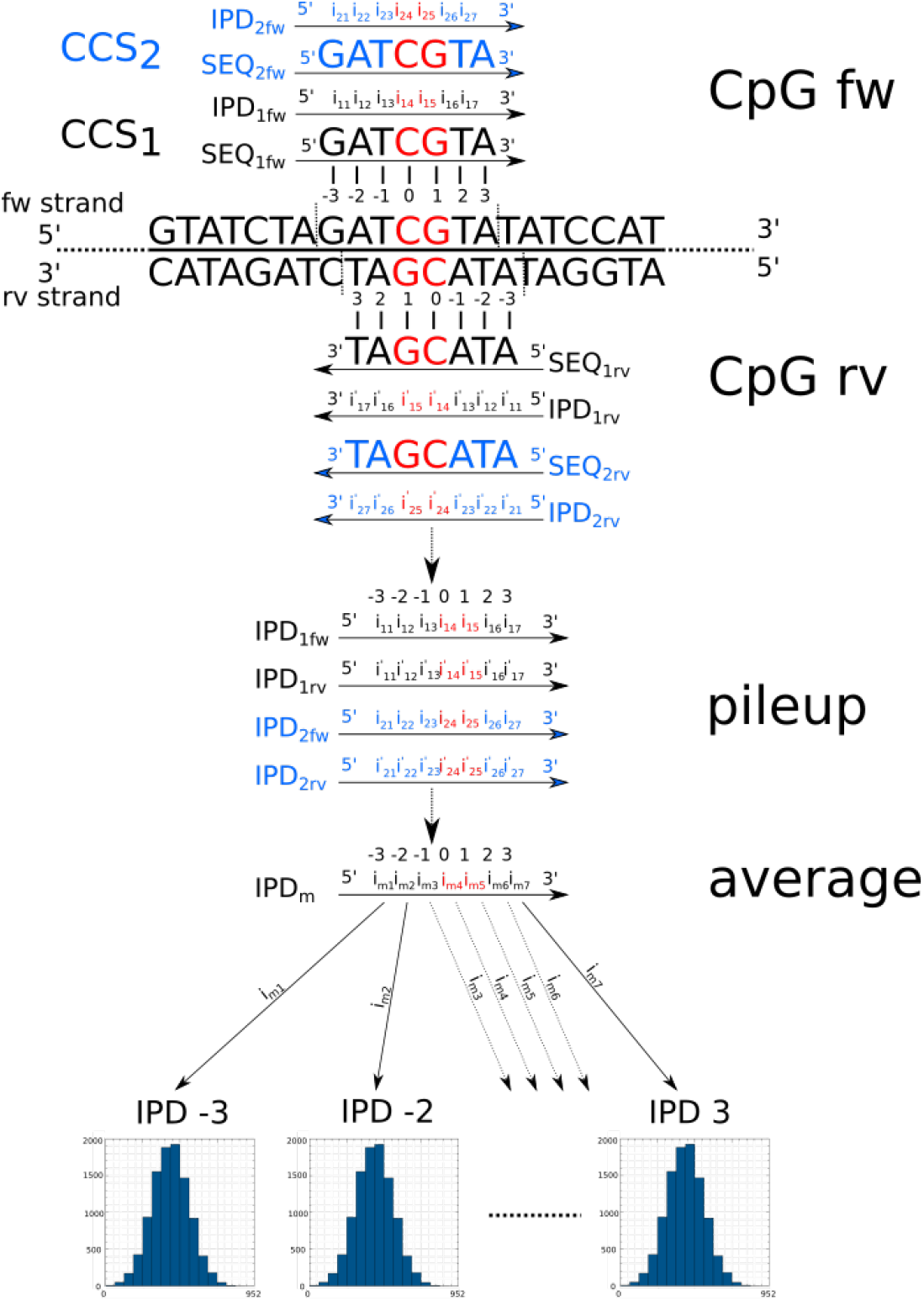
Training of a kinetic signal model: the different steps of the training of a kinetic signal model are displayed. From top to bottom: 1) CCSs mapping over a CpG window - here the window is of size 7bp - are considered, 2) the sub-vector of kinetic values - the IPD only are displayed for clarity - overlapping the CpG window are extracted, 3) a pileup of kinetic vectors is constructed, 4) the strand-aggregated average per-position kinetics are computed from the pileup and 5) each value is stored in the corresponding per-position histogram.

**Fig. S6.**
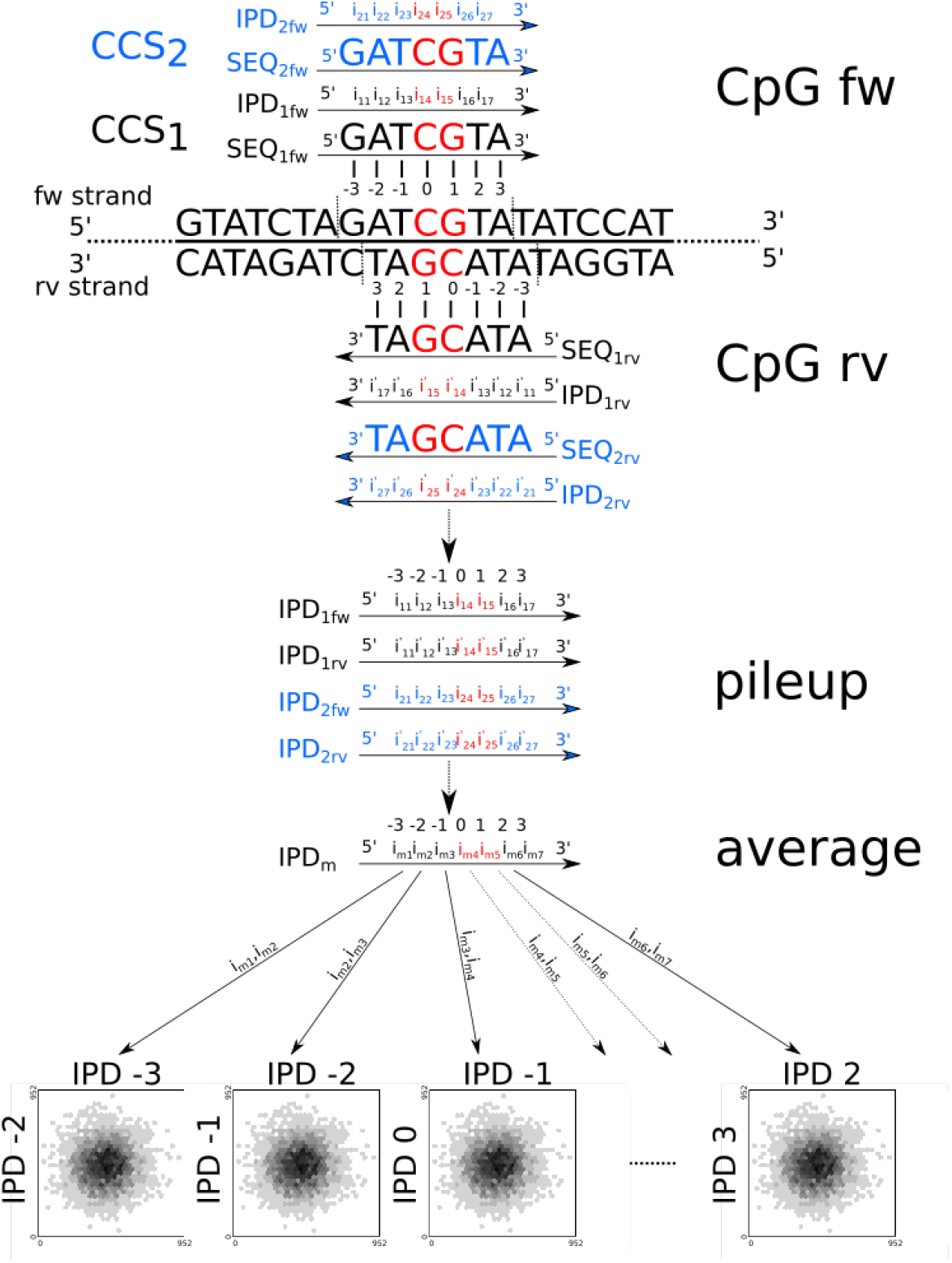
Training of a DP kinetic signal model: the different steps of the training of a kinetic signal model are displayed. From top to bottom: 1) CCSs mapping over a CpG window - here the window is of size 7bp - are considered, 2) the sub-vector of kinetic values - the IPD only are displayed for clarity - overlapping the CpG window are extracted, 3) a pileup of kinetic vectors is constructed, 4) the strand-aggregated average per-position kinetics are computed from the pileup and 5) each pair of neighboring values with index *x, y* ∈ *D*_*DP*_ is fed in thecorresponding per-position histogram.

**Fig. S7.**
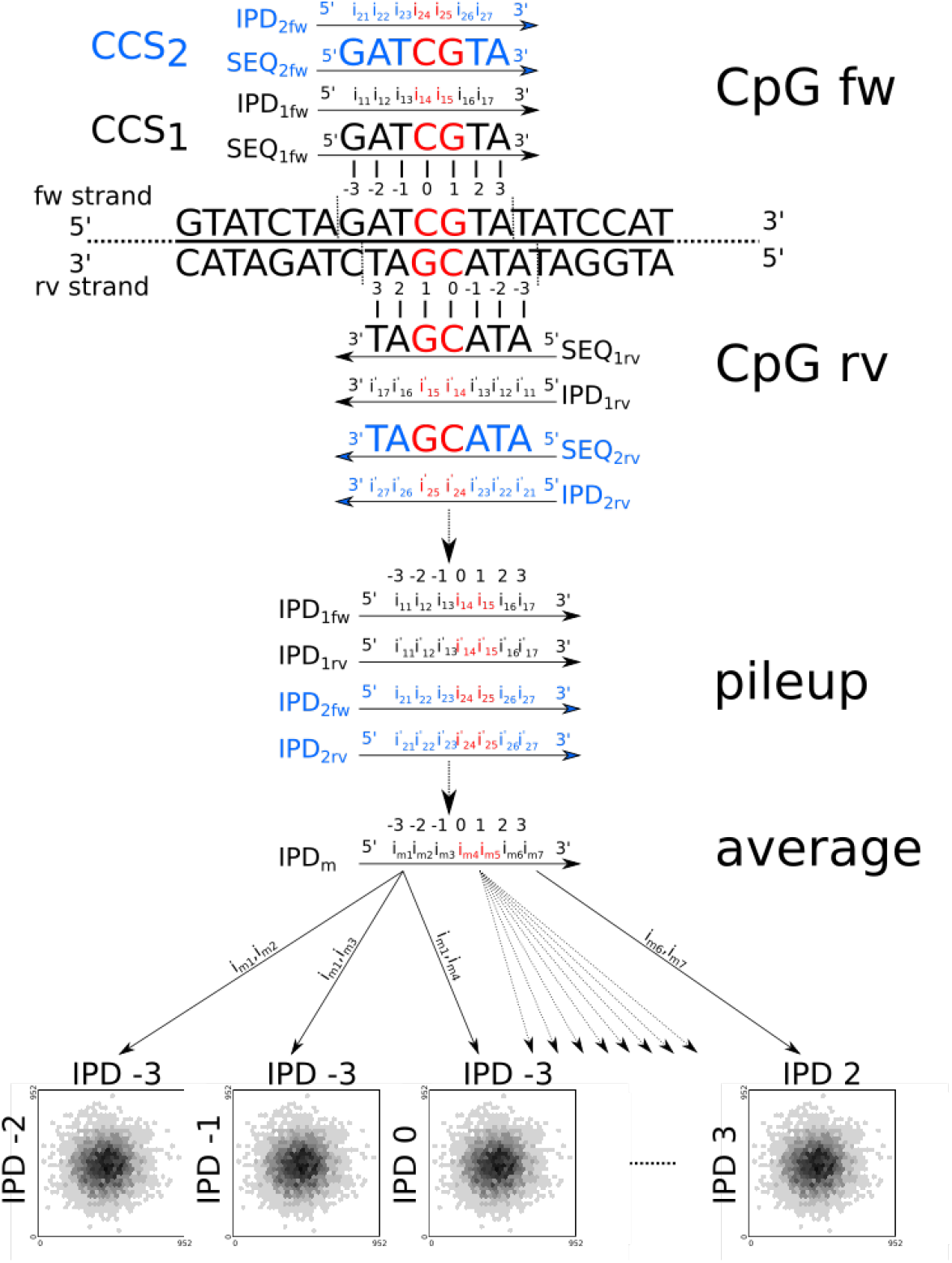
Training of a PW kinetic signal model: the different steps of the training of a kinetic signal model are displayed. From top to bottom: 1) CCSs mapping over a CpG window - here the window is of size 7bp - are considered, 2) the sub-vector of kinetic values - the IPD only are displayed for clarity - overlapping the CpG window are extracted, 3) a pileup of kinetic vectors is constructed, 4) the strand-aggregated average per-position kinetics are computed from the pileup and 5) each pair of neighboring values with index *x, y* ∈ *D*_*P W*_ is fed in the corresponding per-position histogram.

